# Genomic signature of experimental adaptation of *Staphylococcus aureus* to a natural combination of insect antimicrobial peptides

**DOI:** 10.1101/194738

**Authors:** Olga Makarova, Paul Johnston, Alexandro Rodriguez-Rojas, Baydaa el-Shazely, Javier Moreno Morales, Jens Rolff

**Affiliations:** Evolutionary Biology, Institut für Biologie, Freie Universität Berlin, Berlin, Germany; Institute for Animal Hygiene and Environmental Health, Centre for Infection Medicine, Freie Universität Berlin, Berlin, Germany; Berlin Center for Genomics in Biodiversity Research, Berlin, Germany; Leibniz-Institute of Freshwater Ecology and Inland Fisheries (IGB), Berlin, Germany; Zoology Department, Alexandria University, Egyp; Berlin-Brandenburg Institute of Advanced Biodiversity Research (BBIB), Berlin, Germany

## Abstract

Antimicrobial peptides are highly conserved immune effectors across the tree of life and are employed as combinations. In the beetle *Tenebrio molitor*, a defensin and a coleoptericin are highly expressed *in vivo* after inoculation with *S. aureus*. The defensin displays strong *in vitro* activity but no survival benefit *in vivo*. The coleoptericin provides a survival benefit in vivo, but no activity in vitro. To investigate this paradox we experimentally evolved *S. aureus* to increased resistance against the defensin and a combination of the defensin and coleoptericin. Genome re-sequencing showed that resistance was associated with mutations in either the *ytr* or *nsa* operons, in both AMP treatments. Strains with these mutations show longer lag phases, slower Vmax and *nsa* mutants reach lower final population sizes. Mutations in *rpoB* were showed a further increase in the lag phase in *nsa* mutants but not in *ytr* mutants. In contrast, final MICs do not segregate by mutation. All resistant lines display AMP but not antibiotic cross-resistance. Costly resistance against AMPs readily evolves for an individual AMP as well as a naturally occurring combination *in vitro* and provides broad protection against AMPs. Such non-specific resistance could result in strong selection on host immune systems that rely on cocktails of AMPs.

## Introduction

In antibiotic treatments, often single drugs are successfully used to clear infections. Yet, in innate immune systems, infections typically result in the expression and release of cocktails of antimicrobial peptides ^1–3^, even though individual antimicrobial peptides can be very potent ^4^. Possible evolutionary explanations are physiological cost savings, for example, if immune effectors synergize and hence reduce the total amount of effectors required for clearance. Other possible explanations are that resistance evolution has a lower probability if bacteria are under selection from multiple antimicrobials ^5^, or a high probability of mixed infections.

Bacteria have evolved a number of resistance mechanisms that provide protection against ubiquitous AMPs. As most AMPs target the cell envelope and are often cationic, most described resistance mechanisms are related to changes in the cell wall such as altering the net cell surface charge ^6^. In the case of *S. aureus*, the antimicrobial peptide sensing system GraRS ^7^ regulates the *dlt* operon, which controls D-alanylation of the wall teichoic acids ^8^ and *mprF* expression, which is responsible for peptidoglycan lysinylation^6^ as well as the Bce-type ABC transporter vreFG which confers broad-spectrum AMP resistance. Two additional Bce-type ABC transporters, BraDE and vraDE, are under the control of the nisin susceptibility-associated two-component system NsaSR (also known as BceS/BceR ^9^and BraS/BraR ^10^)

In a recent experimental evolution study investigating *S. aureus* resistance evolution against three different AMPs from different organisms (melittin, pexiganan and iseganan ^11^), we found a range of mutations associated with resistance ^12^. These mutations did not segregate by AMP.

An interesting observation is that during an infection antimicrobial peptides are expressed that have no known activity against the agent of infection. This is surprising, given that insects for example have different receptors that can distinguish between classes of infectious microbes such as fungi or bacteria ^13^ and energetic costs of protein synthesis are considered to be high^3^. Moreover, recent studies, for example in bumble bees, have also shown pathogen strain specific responses on the level of the transcriptome ^14^. In the mealworm beetle *Tenebrio molitor* experimental infection with *S. aureus* induces the expression of at least ten antimicrobial peptides for a week ^3^. A proteomic analysis shows that the majority of these inducible antimicrobial peptides remain elevated in the haemolymph after three weeks ^15^. While some of them such as Tenecin 1, a defensin, show high activity against *S. aureus in vitro*, other abundant AMPs such as Tenecin 2, a coleoptericin, display no activity against *S. aureus* ^16^. This is contrasted by studies using a gene knock-down approach *in vivo* ^17^. While the knock-down of *tenecin 1* did not change host survival, the knock-down of *tenecin 2* led to highly increased mortality of *T. molitor* 3 days after *S. aureus* infection. This clearly indicates that Tenecin 2 has an unknown role or activity *in vivo*.

Here we explore antimicrobial peptide resistance evolution *in vitro* against a pair of naturally co-expressed AMPs in *T. molitor*. Using an experimental evolution protocol, we selected *S. aureus* for resistance against the *T. molitor* defensin Tenecin 1, either alone or in combination with Tenecin 2. We then investigated if resistance evolution results in fitness costs and if the presence of Tenecin 2 changes the outcome of resistance evolution against the potent Tenecin 1. The resulting strains were re-sequenced to identify mutations associated with AMP resistance and to study the degree of parallel evolution.

## Materials and Methods

### Bacterial strains and culturing conditions

*Staphylococcus aureus* strain SH1000 and *Escherichia coli* strain MG1655 were used in the experiments. Bacterial cultures were grown in non-cation adjusted Mueller Hinton Broth (MHB) (Panreac Applichem GmbH) at 37°C with mild shaking and plated on Mueller Hinton Agar (MHA), unless stated otherwise.

### Antimicrobial peptides

Mealworm *Tenebrio molitor* antimicrobial peptides Tenecin 1(VTCDILSVEAKGVKLNDAACAAHCLFRGRSGGYCNGKRVCVCR) and Tenecin 2 (SLQPGAPSFPGAPQQNGGWSVNPSVGRDERGNTRTNVEVQHKGQDHDFNAGWGKVIKGKEKGSPTWHVGGSFRF) were chemically synthesised by Peptide Protein Research Ltd (Funtley, UK). To avoid multiple freeze-thaw cycles and prevent binding of the peptides to the vials during storage, peptides were re-suspended in sterile water to the final peptide concentration of 5 mg/ml and glycerol concentration 50% and stored at -20°C in sterile glass vials, which were pre-treated with “Piranha” solution (3 parts of concentrated sulfuric acid and 1 part of 30% hydrogen peroxide solution).

### Selection experiment

Prior to selection, bacteria were pre-adapted to the experimental conditions (following ^18^,^19^). For this, three randomly selected clones of *S. aureus* strain SH1000 were picked from a Tryptic Soy agar plate, inoculated individually into 10 ml MHB and incubated overnight with shaking at 37°C. To mimic experimental conditions, the cultures were then diluted 1:1000 and incubated at 37°C without agitation in 50 ml polypropylene Falcon tubes containing 3.7 ml MHB. The specific volume of MHB used for pre-adaptation was calculated to ensure the same surface-area-to-volume ratio as in 96-well plates that were used in the evolution experiment. Pre-adaptation was carried out as described above by serial passage every 24 hours (to allow for approximately 36 doublings) for 8 days, with daily measurements of optical density at 600 nm, contamination checks by plating out on MHA and cryopreservation of culture aliquots at -80°C in 12% glycerol solution. One “ancestor” line was used to establish the initial MIC value for Tenecin 1.

For the selection protocol, five independent parallel selection lines (numbered 1 to 5) were founded by plating the pre-adapted “ancestor” line on MHA and isolating five random colonies. The experiment was performed at 37°C without shaking in a microplate reader (Synergy 2, Biotek). We used flat bottom polypropylene non-binding 96-well plates (Greiner Bio-One GmbH, Germany) to avoid attachment of the peptides to the plastic surfaces covered with clear polystyrene lids with condensation rings (Greiner Bio-One GmbH, Germany). The plates were filled with MHB, with the total volume of 200 μl per well. Growth curves were generated by taking measurements of OD_600_ every 30 minutes (preceded by a brief shaking for 10 seconds) for 23 hours. For each of the five replicate lines there were two experimental conditions – Tenecin 1 or a combination of Tenecin 1 + Tenecin 2, as well as four controls - negative control (culture medium control), two glycerol controls - for each AMP treatment - to account for the increasing concentrations of glycerol at higher concentrations of peptides, in which they were stored, and a non-selected control. The serial passage started at 1/2 x MIC, which corresponded to 4 μg/ml for Tenecin 1 and 8 μg/ml for Tenecin 2. To inoculate the treatment wells and the respective control wells, overnight cultures of the five replicate lines were diluted 1:100 and sub-cultured until OD_600_ 0.5 (corresponding to 1x10^8 cfu/ml), then 10 μl of these cultures were inoculated into each treatment and control (except NTC) wells resulting in the final total volume of 200 μl and bacterial density of approximately 1x10^6 cfu per well. If the OD_600_ values in the AMP treatment wells were at least half that of the respective non-selected controls after 23 hours of incubation, 2 μl (1:100) from each treatment and control well were transferred using a multichannel pipette to a fresh 96 well plate and the concentration of AMPs was doubled. Twenty μl of the remaining cultures were added to 180 μl of sterile 0.9% NaCl and then serially diluted (typically from 10^-1 to 10^-5) and checked for contamination and viable counts by plating 5 μl of each dilution on MH-agar using the drop plate method and incubating the plates overnight at 30°C. Glycerol was added to the rest of the cultures to the final concentration of 12% and the plates were stored at -80°C. The selection experiment continued for 7 passages, in each of which the concentration of AMPs was doubled reaching 256 μg/ml for Tenecin 1 and 512 μg/ml Tenecin 2 on the last day of the experiment (day 7). When the volume of added peptide started to exceed 5% of the total volume of medium per well, 2-fold concentrated MHB was used to prepare stock solutions of the desired concentration to alleviate the possible effects of nutrients depletion. The resulting resistant lines were used for subsequent assays (MIC, growth curves) and sequencing both as populations and colonies.

### Antimicrobial susceptibility testing

Minimal inhibitory concentration (MIC) was determined using a broth micro-dilution method ^20^. Briefly, 5 μl (1x10^5 cfu/ml) of the mid-exponential phase bacterial culture diluted 1:100 were inoculated into the wells of polypropylene V-bottom 96-well plates (lids with condensation rings 656171, both from Greiner Bio-One GmbH, Germany, Germany) containing two-fold serial dilutions of AMPs or antibiotics in the total volume of 100 μl MHB per well. Each assay was performed in triplicate. The plates were incubated at 37°C in a humidity chamber. The MIC was defined as the lowest concentration that inhibited visible bacterial growth after 24 hours of incubation. This standard method was used to determine the initial MIC of Tenecin 1 for the pre-adapted “ancestor” line and mutant lines and antibiotics. Because Tenecin 2 is mostly active against gram-negative bacteria, *Escherichia coli* strain MG1655 was used to determine the activity of this peptide.

To determine the MICs of Tenecins immediately after the selection experiment, we scaled-down the assays to the total volume of 40 μl per well because of the high number of bacterial lines and the limited amount of the antimicrobial peptide available. For this, we used flat-bottom 384-well polypropylene plates (Greiner Bio-One GmbH, Germany) and clear polystyrene lids (Greiner Bio-One GmbH, Germany). We determined MIC as the lowest concentration at which OD_600_ readings were indistinguishable from those of the NTC wells. As a method control, we compared the MIC determined using the standard and the small-volume protocol, and found no differences. The MICs were determined for populations and individual colonies derived from the selection lines.

### Growth curves

Growth curve assays were performed by monitoring the changes in turbidity at OD_600_ of the selected mutant lines, non-selected controls and ancestor in un-supplemented MHB using a microtitre plate reader. For this, bacterial lines (populations and colonies) were grown until OD_600_ 0.5, diluted 1:10 and 20 μl of the resulting cell suspension were inoculated into 180 μl MHB. Each assay had four replicates. The measurements were taken at 20 minutes intervals during 16 hours of incubation at 37°C inside the microtitre plate reader, with 10 seconds shaking before each reading. Growth parameters such as final and maximum OD, Vmax and lag time were calculated with Gen5 software (Biotek).

### DNA isolation

Genomic DNA for whole genome sequencing was isolated using GeneMATRIX Bacterial and Yeast genomic DNA purification kit (Roboklon, Germany) following manufacturer’s instructions. Four μl of 10 mg/ml freshly prepared lysozyme and lysostaphin (both from Sigma) each were added into bacterial lysate. The DNA quantity and quality were estimated by measuring the optical density at A260/280 using the Nanodrop spectrophotometer (Thermo Scientific) and agarose gel electrophoresis.

### Genome re-sequencing

TruSeq DNA PCR-free libraries were constructed according to the manufacturers instructions and sequenced for 600 cycles using a MiSeq at the Berlin Center for Genomics in Biodiversity Research. Sequence data are available from the NCBI SRA under BioSample accession xxxxxxxx.

The genetic differences between strain SH1000 and other members of the 8325 lineage have been described using array-based resequencing ^21^, and de novo genome sequencing^22^. The differences comprise: the excision of three prophages from 8325 (Φ11, 12, 13), 13 single-nucleotide polymorphisms (SNPs; two synonymous, 11 nonsynonymous), a 63-bp deletion in the spa-sarS intergenic region, and an 11-bp deletion in rsbU ^22^. To account for these differences we first assembled reads from SH1000 using SPAdes ^23^, and used the resulting contigs to correct the three phage excision sites in the 8325 reference genome. SH1000 reads were then mapped to the resulting sequence and bcftools consensus ^24^ was used to correct the remaining 13 SNPs and two indels.

The haploid variant calling pipeline snippy^25^ was used to identify mutations in the selection lines. Snippy uses bwa^24^ to align reads to the reference genome and identifies variants in the resulting alignments using FreeBayes (Garrison and Marth 2012). All variants were independently verified using a second computational pipeline, breseq ^26^.

### Statistical analyses

Statistical analyses were performed using R version 3.4.1. Growth parameters (Vmax, duration of lag phase, and final OD600) were analysed by using the nlme package ^27^ to fit linear mixed-effects models specifying line as a random effect. We also used visreg ^28^ and Ismeans ^29^. Permutational analyses of Jaccard distance were performed using the vegan package^30^.

## Results

### Resistance evolves readily

Over the course of selection all replicates of both treatments (single and combination of AMPs) readily evolved a high level of resistance. All lines were able to grow in the presence of 256 μg/ml Tenecin 1 and 512 μg/ml Tenecin 2, resulting in an increase in MIC for Tenecin 1 (T=-9.98, df=35, p=<0.0001, Fig. 1A).

**Fig 1.**
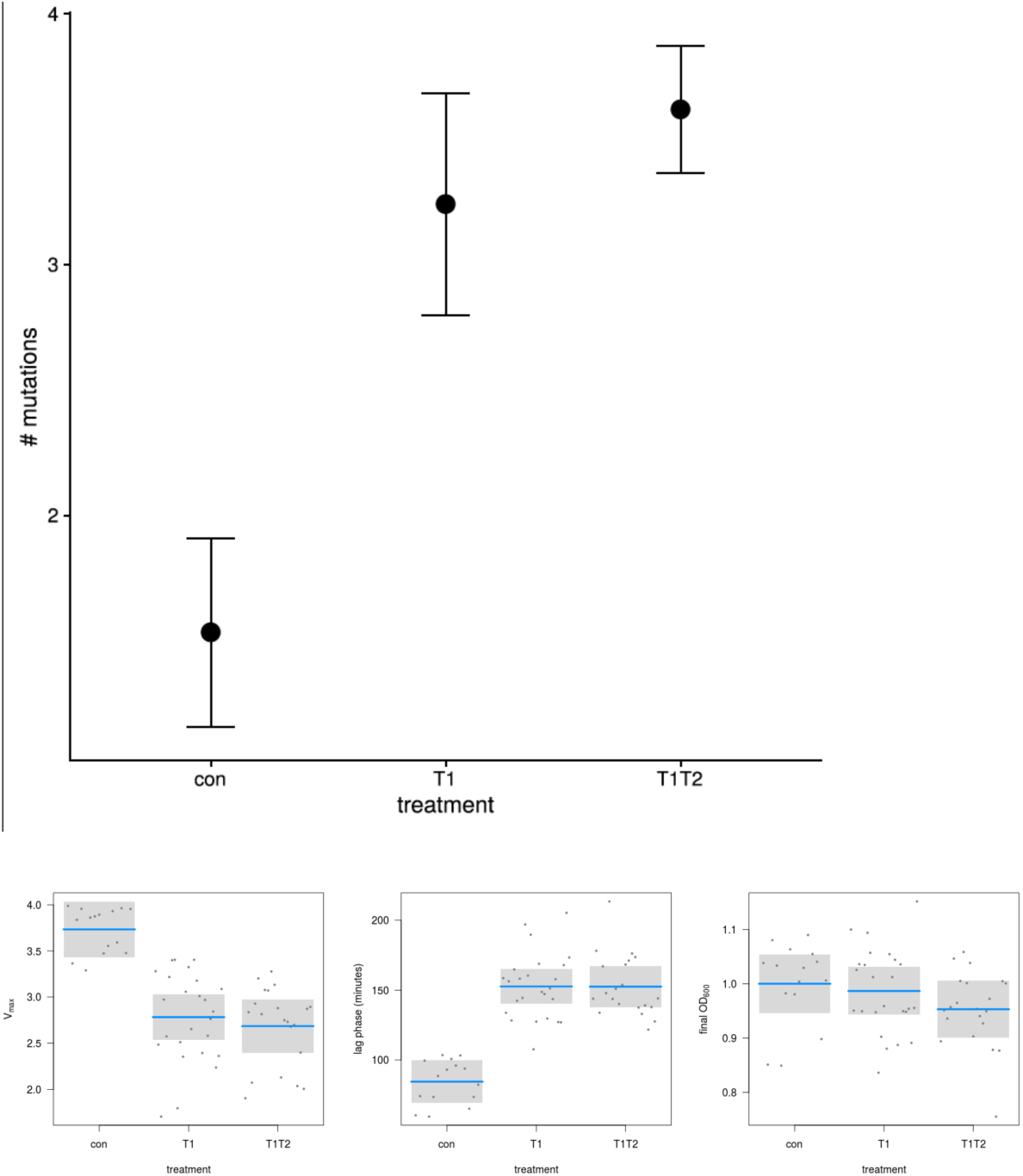
(A) Increase in MIC in *S. aureus* selected under either Tenecin (Defensin) or combined Tenecin 1 and Tenecin 2 (Coleopterecin) (mean + 95% CI). (B) Growth parameters of experimentally evolved AMP-resistant *S. aureus* in relation to treatment (Vmax, lag phase, maximal OD).

### Reduced fitness in AMP-resistant mutants

When comparing bacterial growth curves of the mutants with those of non-selected controls and ancestors, we found consistently slower growth rates in the exponential phase for both Tenecin 1 (T = -6.8, df = 56, p = <0.0001, Fig. 1B) and Tenecin 1+Tenecin 2-selected lines (T = -7.17, df = 56, p = <0.0001, Fig. 1B) and also extended lag phases (Tenecin 1: T = 9.4, df = 56, p = <0.0001, Tenecin 1+Tenecin 2: T = 9.1, df = 56, p = <0.0001, Fig. 1B), Tenecin 1 and Tenecin 1+Tenecin 2 lines did not differ (T = -0.67, df = 56, p = 0.5042, Fig. 1AB). This suggests that AMP resistance incurs fitness costs irrespective of the type of selection.

### Cross-Resistance to other AMPs but not antibiotics

We tested for cross-resistance against commercially available AMPs, colistin, pexiganan and melittin. We found 2-8-fold cross-resistance of all AMP-selected strains against the AMPs Mellitin, Pexiganan and Colistin (table S1). The selected strains showed no cross-resistance against Vancomycin, a drug of last resort for staphylococcal infections with multiple resistances. There was no relationship between the AMP cross-resistance and the resistance mutations (see table S2). We also tested the evolved strains against a panel of eight antibiotics (see table S2) but detected no cross-resistance.

### Genome re-sequencing reveals mutations in a limited number of loci

Whole genome sequencing of the selected mutants and the respective controls (at the population and single colony levels) showed differences both between treatments and between replicate lines within treatments. In each resistant strain one mutation was identified in either *ytrA*, *ytrB*, *nsaS*, or *nsaR* gene, all of which are known to be involved in envelope stress tolerance (see also table S3 for a full list of mutations). In case of the ytrA mutations, there were examples of missense, frameshifts, and stop gains. For *ytrB, nsaS, nsaR, rpoB*, and *rpoC* mostly missense mutations were found. Interestingly, we found the same *ytrA* stop-gain mutation (c.14T>A p.Leu5*) in at least 11 strains, which has previously been described for melittin-selected *S. aureus* lines (Johnston et al 2016). Additionally we also identified an *nsaS* missesnse mutation (A_208_E) in 4 strains which has been shown to be responsible for nisin resistance in SH1000^31^, the same strain as used here. The final MICs achieved did not segregate by mutation (T=0.25, df=32, p=0.8, Fig 2).

**Fig 2.**
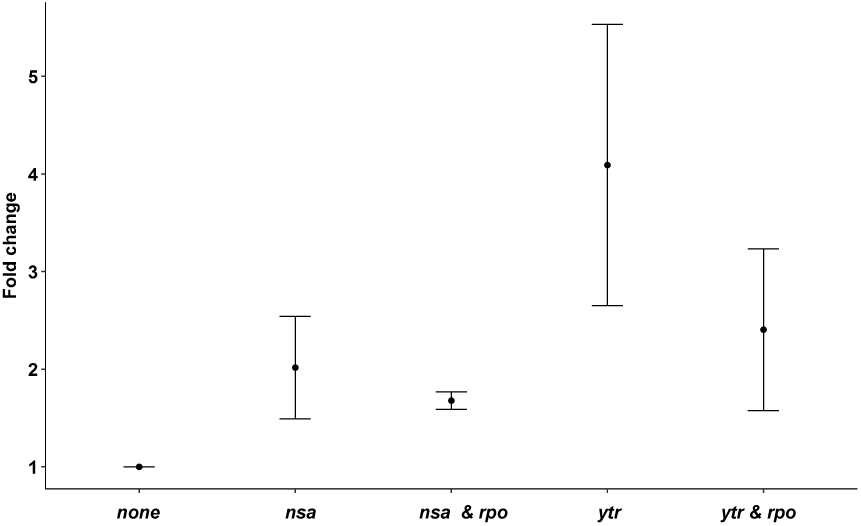
Fold change in MICs for resistant strains separated by mutations compared to procedural controls.

### Reduced fitness in relation to resistance mutations

Fitness reduction, as measured by the increased lag phase, did not segregate by mutation (T = 0.93, df = 56, p = 0.3576, Fig. 3A). Vmax differed between *nsa* and *ytr* mutations, with *nsa* mutants showing a lower growth rate than *ytr* mutants (T = 2.7, df = 56, p = 0.0092) and both mutants grow significantly slower than the controls (*ytr*: T = - 6.46, df = 56, p = <0.0001, *nsa*: T = -8.16, df = 56, p = <0.0001, Fig.3B). In the presence of a second mutation *rpo*, *nsa* mutants show a further extension of the lag phase (T = 4.21, df = 54, p = 1e-04, Fig.3B).

**Fig 3.**
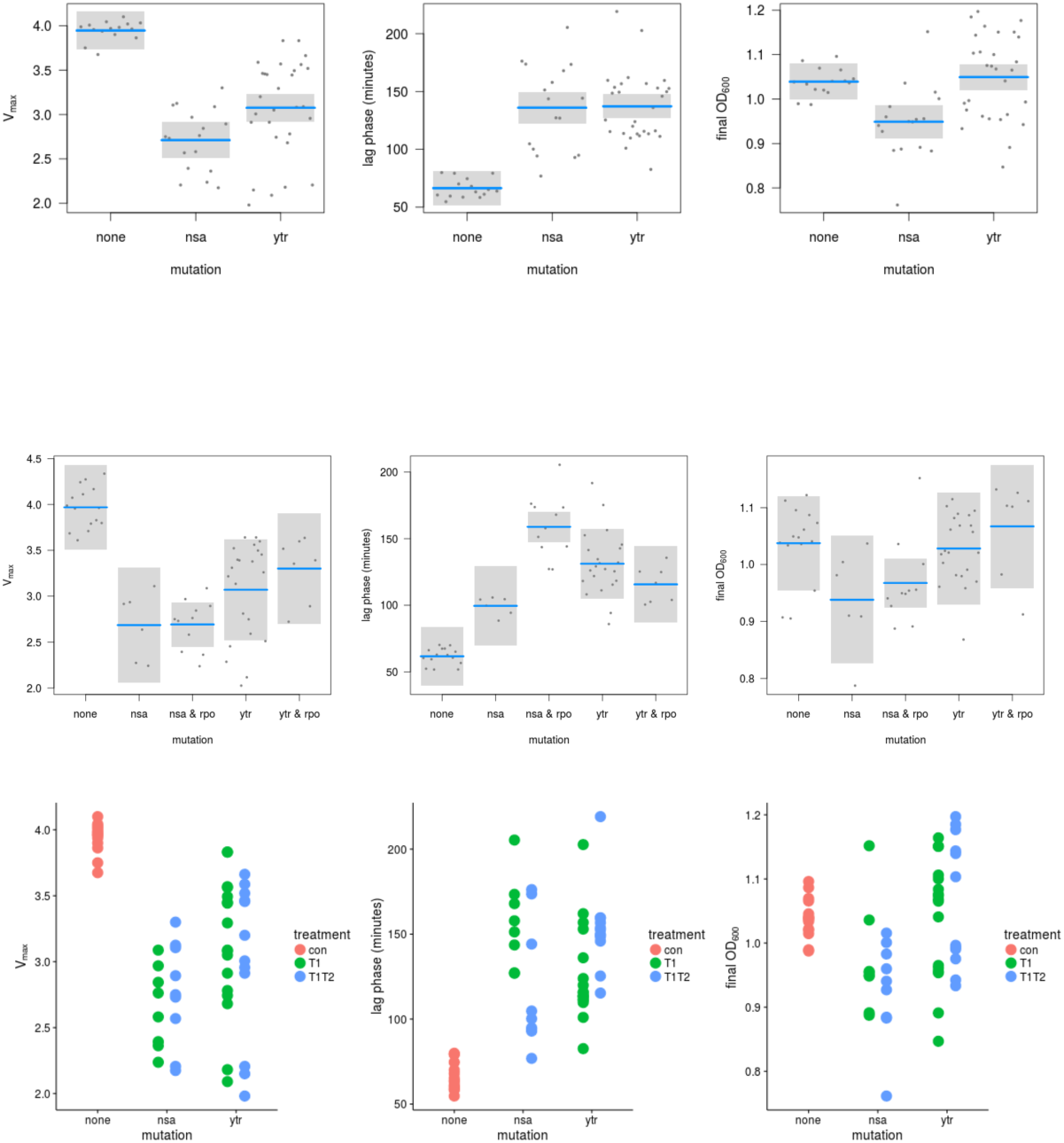
Fitness costs over mutation/operon (a) and in the presence or absences of a second mutation (b). Fitness of mutations (c) in relation to the selective environment.

### Parallel evolution

Using the Jaccard distance to calculate the degree of parallel evolution, we did not find evidence for parallel evolution at the operon level. Selection treatment (T1 or T1T2) did not affect the mean proportion of shared mutated operons (permutational analysis of multivariate homogeneity of group dispersion F = 0.1925, p = 0.625) or the mean number of shared mutations (permutational multivariate analysis of Jaccard distance matrix, F = 1.063, p = 0.347, Fig. 4). Yet, as reported above, all AMP selected lines showed either a mutation in *nsa* or in *ytr* operons, but never in both (table 2).

**Fig 4.**
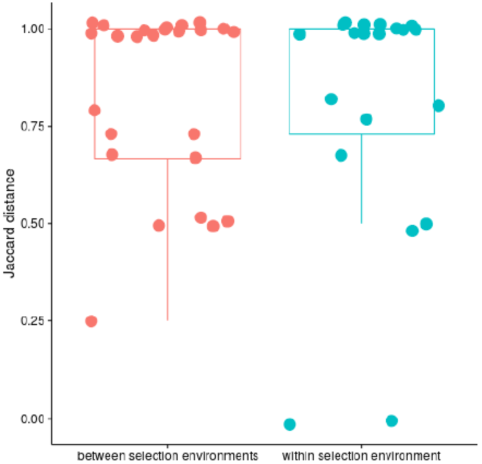
Parallel evolution as assessed by Jaccard Distance for operons (1=no shared evolution).

## Discussion

Our study was one of the first to explore the evolution of resistance to AMPs that are part of the same immune system. We find that all populations under antimicrobial peptide selection evolve resistance quickly and to a similar degree irrespective of the presence of one or two AMPs. In all cases resistant strains possessed a mutation in either in the *ytr* or *nsa* operons. Mutations in these operons were not found together.

We have previously shown that evolution of resistance towards the AMP melittin in *S. aureus* JLA513 is associated with nonsense mutations in *ytrA*. Here we find that the most frequent *ytrA* mutation is identical to the stop-gain mutation described in *S. aureus* JLA513^12^. In *S. aureus*, the *ytr* operon is induced by cationic AMPs ^32^ and encodes a putative GntR-family repressor (YtrA) as well as two ABC transporters. In *Bacillus subtilis*, *ytr* repression requires *ytrA* and null mutations lead to constitutive *ytr* expression. This raises the possibility that the *ytr* mutations observed here may mediate Tenecin 1 resistance via derepression of *ytr*.

The nisin susceptibility-associated (*nsa*) two-component system was independently discovered three times^9^,^10^,^31^ as being responsible for resistance to nisin and bacitracin. In the presence of it’s substrate NsaSR activates transcription of the Bce-AB type ABC transporters BraDE and VraDE which are involved in sensing and detoxification respectively10. Strikingly, in 4 strains we identified the same *nsaS* mutation (A_208_E) which was shown to confer increased nisin resistance in SH1000^31^.

In general antimicrobial resistance is costly ^33^. While this is a consistent finding for antibiotics, the patterns for AMPs are less clear ^19^,^34^. Here we find clear evidence for costly resistance as measured in growth rate and lag phase. With the exception of the double mutation *nsa/rpoB*, which displays the longest lag phase the costs in our experiments do not segregate by mutation. *rpoB* mutations in AMP-resistant *S. aureus* have been found before ^35^. At the moment it is not clear why the more costly double-mutation evolves, given that there is no difference in MICs between *nsa* and *nsa/rpoB* mutants. Limited costs in *S. aureus* selected for resistance against the human AMP LL-37 were reported^35^, but the same study found reduced growth rates in AMP-resistant strains. Antibiotic resistance evolution resulting in extended lag phases has been frequently reported (e.g. ^36^) and also has been proposed as a mechanism explaining antibiotic tolerance ^37^ even in the absence of *bona fide* resistance.

## Lack of parallel evolution

While we do find strong evidence for parallel evolution at the level of the resistance phenotype, all populations under selection evolve resistance at a comparable speed to a similar level (MIC), we do not find evidence for parallel evolution at the level of the operons and treatments. This contrasts with findings reported in *Pseudomonas fluorescens* under antibiotic selection^38^, which found a higher degree of parallel evolution, albeit in more complex environments. A low level of parallelism has been observed in other studies of bacterial antibiotic resistance evolution^36^,^39^. While our assessment is based on the Jaccard-Distance, it is noteworthy that all strains have one of two resistance-associated mutations. This is much lower than the variation observed in a previous study in *S. aureus* against a panel of antimicrobial peptides originating from different organisms ^12^.

### Cross-resistance

Despite the limited number of resistance mutations, both our experimentally evolved Tenecin 1 and Tenecin 1+Tenecin 2-selected strains, display cross-resistance against colistin, pexiganan and melittin. All of these antimicrobial peptides have a different origin (from *Paenibacillus polymixa*, the African clawed frog and the honey bee, respectively) and belong to distinct AMP families. This underlines that AMP resistance mechanisms can be relatively nonspecific. In gram-positive bacteria resistance is mostly achieved by modifications to the physico-chemical properties of the cell envelope ^6^.

Cross-resistance between AMPs has been reported before^19^,^34^ and might constitute a risk for the application of AMPs in medical treatments^40^. In the context of natural immune defenses it is likely that the resistance evolution to one AMP possibly results in resistance also against AMP cocktails. This could constitute a strong selection pressure for those innate immune systems that are strongly dependent on AMPs. Finally, while we did not find indications for cross-resistance against antibiotics, cross-resistance against human AMPs and the peptide antibiotic daptomycin, which targets the cell membrane, has been reported in MRSA ^41^. It hence seems possible that resistance against AMPs comes with the risk of cross-resistance against new potential antibiotics such as teixobactin ^42^ that target the cell surface.

## Acknowledgements and funding information

We thank Vitali Laba for support in the lab. This work was supported by the European Research Council (grant 260986, OM, PJ, JR) the Deutsche Forschungsgemeinschaft (SFB 973, C5, ARR, JR) the German Academic exchange service (personal stipend to BES) and ERASMUS (personal stipend to JMM). Accession number for Genome Data: PRJNA399645.

